# The influence of thermal extremes on coral reef fish behaviour in the Persian Gulf

**DOI:** 10.1101/654624

**Authors:** Daniele D’Agostino, John A. Burt, Reader Tom, Grace O. Vaughan, Ben B. Chapman, Santinelli Veronica, Geórgenes H. Cavalcante, David A. Feary

## Abstract

Despite increasing environmental variability within marine ecosystems, little is known about how coral reef fish species will cope with future climate scenarios. The Arabian/Persian Gulf is an extreme environment, providing an opportunity to study fish behaviour on reefs with seasonal temperature ranges which include both values above the mortality threshold of Indo-Pacific reef fish, and values below the optimum temperature for growth. Summer temperatures in the Gulf are comparable to those predicted for the tropical ocean by 2090-2099. Using field observations in winter, spring and summer, and laboratory experiments, we examined the foraging activity, distance from refugia and resting time of *Pomacentrus trichrourus* (pale-tail damselfish). Observations of fish behaviour in natural conditions showed that individuals substantially reduced distance from refugia and feeding rate and increased resting time at sub-optimal environmental temperatures in winter (average SST = 21°C) and summer (average SST = 34°C), while showing high movement and feeding activity in spring (average SST= 27°C). Diet was dominated by plankton in winter and spring, while fish used both plankton and benthic trophic resources in summer. These findings were corroborated under laboratory conditions: in a replicated aquarium experiment, time away from refugia and activity were significantly higher at 28°C (i.e., spring temperature conditions) compared to 21 °C (i.e., winter temperature conditions). Our findings suggest that *P. trichrourus* may have adapted to the Arabian/Persian Gulf environment by downregulating costly activity during winter and summer and upregulating activity and increasing energy stores in spring. Such adaptive behavioural plasticity may be an important factor in the persistence of populations within increasing environmentally variable coral reef ecosystems.

## Introduction

Over the last century sustained climate change has resulted in an average global sea surface temperature increase of 0.6°C, with predictions of further 4.0°C rises by 2100 (Collins et al. 2013; IPCC 2014). As the planet warms, the occurrence of extreme weather events will also increase (IPCC 2012) and, as the world’s oceans absorb 93% of excess heat trapped by greenhouse gases (Laffoley and Baxter 2016), this will have major consequences for the biodiversity, functional structure and productivity of marine ecosystems (Folguera et al. 2011; Doney et al. 2012; Bellard et al. 2012; Thornton et al. 2014; Pecl et al. 2017). Alterations have already being documented in the diversity and assemblage structure in temperate seaweed and seagrass ecosystems (Wernberg et al. 2013; Vergés et al. 2014), coral reefs (Hughes et al. 2018) and costal temperate rocky reefs (Lejeusne et al. 2010; Albouy et al. 2014). Despite this, our understanding of the capacity for marine ecosystems to resist or show resilience to climate change is limited (Côté and Darling 2010).

Coral reefs are particularly susceptible to a range of natural and anthropogenic disturbances that have resulted in degradation and loss of coral on a global scale in recent decades (Wilkinson 1999; Cheal et al. 2017; Hughes et al. 2018). While these disturbances are clearly significant for the corals that serve as the biogenic engineers of these ecosystems, diverse and ecologically important reef-associated fish assemblages also often exhibit dramatic changes in composition and abundance with increasing environmental variation (Day et al. 2018; Richardson et al. 2018; Gordon et al. 2018). Indeed, in addition to susceptibility to habitat loss (Pratchett et al. 2018), coral reef fishes are also directly threatened by ocean warming due to their narrow thermal tolerance range (Tewksbury et al. 2008; Sunday et al. 2011). Experimental evidence suggests that coral reef fish may already be living close to their upper thermal limits (Rummer et al. 2014) and that population structure may be directly impacted by increased variability in water temperature (Folguera et al. 2011; Pratchett et al. 2015; Rodgers et al. 2018).

One of the primary responses to environmental temperature change is modification of behaviour, with the speed and scope of behavioural adjustment potentially determining survival (Tuomainen and Candolin 2011; Wong and Candolin 2015). Tropical reef fishes, however, have evolved in quite thermally stable environments and may have little ability to adjust their behaviour adaptively (Candolin 2018). Furthermore, as fish metabolism and neurophysiology are directly influenced by temperature (Pörtner and Farrell 2008; Pörtner et al. 2010), environmental temperatures outside an individual’s optimum range may have a pervasive effect on ecological performance and survival via non-adaptive (dysfunctional) modification of behaviour, altering individual feeding rate, risk-taking behaviour, activity levels and habitat use (Figueira et al. 2009; Nagelkerken and Munday 2016). Despite this, a growing body of research suggests adaptive behavioural plasticity in response to temperature increases in coral reef fishes, often through temporary reductions in aerobically-costly behaviours (i.e. activity and feeding) associated with increased (short-term) environmental variance (Nowicki et al. 2012; Johansen et al. 2014; Scott et al. 2017; Chase et al. 2018). Nevertheless, how well coral reef fishes will cope with ocean warming scenarios (ICCP 2014), and by what mechanisms, is still generally poorly understood.

One approach to understanding how individuals may behaviourally respond to environmental variance is to study contemporary communities that exist within naturally variable environments. The fauna associated with the high latitude reefs of the southern Arabian/Persian Gulf (hereafter the ‘Gulf’) provide a natural test of the impact of extreme water temperature variation on coral reef fish behaviour (Feary et al. 2010; Burt et al. 2011b). The Gulf experiences the highest annual change in water temperature for coral reefs globally (winter: <15°C, summer: >35°C), with fishes persisting for several months in conditions that would be considered lethal to reef fishes in other parts of the world (Riegl and Purkis 2012; Vaughan et al. 2019). Summer temperatures in the Gulf exceed the upper thermal limits of most tropical reef fishes (Nilsson et al. 2009; Munday et al. 2009; Rummer et al. 2014; Rodgers et al. 2018), while winter temperatures are well below the optimum temperatures reported for reef fish elsewhere in the Indo-Pacific (Eme and Bennett 2008; Figueira and Booth 2010; Nakamura et al. 2013). Indeed, contemporary summer water temperatures in the Gulf are comparable to those predicted for tropical oceans at the end of this century (Riegl and Purkis 2012; IPCC 2014), while winter conditions can be so severe as to induce cold water coral bleaching (Shinn 1976; Coles et al. 1991).

Using a combination of *in situ* field observations and *ex situ* experimental manipulations, this study examined the influence of extreme thermal variability on reef fish behaviour in the southern Gulf. Using the locally abundant *Pomacentrus trichrourus* (pale-tail damselfish; Günther, 1867) we performed field-based observations of behaviour across three thermally distinct seasons separated by a 13 °C sea surface temperature (SST) range (winter: SST = 21 °C, spring: 27 °C, and summer: 34 °C), monitoring individual distances from refugia, resting times, feeding rate and diet across three sites in the southern Gulf. We predicted that fish would respond to low and high water temperature extremes by modifying their behaviour (i.e., exhibit behavioural plasticity) to different temperature conditions. Specifically, we hypothesize that fish would minimize aerobically-costly behaviours (i.e., reduced distance from refugia and feeding rate, whilst increasing resting time) when exposed to sub-optimal temperatures in winter and summer, while compensating for energy losses and building up energetic stores by increasing activity and feeding in spring. We also hypothesized that *P. trichrourus* would exhibit flexibility in diet by shifting from a predominately planktonic diet during optimal condition to a mixed planktonic and benthic diet when exposed to suboptimal temperature. By restricting foraging movements in suboptimal conditions, fish may have to become more generalist in their feeding habits, exploiting more easily accessible resources. Finally, we performed an aquarium-based experiment exposing *P. trichrourus* to temperatures typical of winter (21 °C) and spring (28 °C) in order to examine the hypothesis that variation in behaviour observed in the field was directly mediated by temperature. The results of this study have important implications for understanding how coral reef fishes may use behavioural modification as a means to respond to the environmental variability that is expected to increase under future climate change.

## Materials and methods

The pale-tail damselfish is a small omnivorous damselfish common in southern Gulf coral communities, often living in association with a single ‘home’ coral head (Allen 1991; Shraim et al. 2017). Within the Gulf this species shows high abundance, a benthic life style with diurnal behaviour, a strong association with the reef, and high territoriality to a small area [territory size of 0.5 to 1.5 m^2^]). It has a small body size (max 6 cm standard length - SL -) and distinct differences in colour pattern between juvenile and adults (Randall 1995; Feary et al. 2010). All individuals within this study were adults (∼4 - 6 cm SL). The optimum temperature for this species is modelled to be between 24.7 °C and 29.3 °C (mean 27.2 °C) (Kaschner et al 2016).

### *In situ* behavioural quantification of *P. trichrourus*

To determine the effect of seasonal changes in temperature on behaviour of *P. trichrourus* in the Gulf, behavioural observations were replicated across three seasons: spring (April/May 2016, average SST = 27°C), summer (August/September 2016, average SST = 34°C), and winter (February/March 2017, average SST = 21°C). Observations were undertaken at three reef sites spaced ca. 40 km apart from one another along the Abu Dhabi coast (United Arab Emirates – southern Gulf): Dhabiya (24°21’55.8″N, 54°06’02.9″E), Saadiyat (24°35’56.4″N, 54°25’17.4″E) and Ras Ghanada (24°50’53.4″N, 54°41’25.1″E) (Fig. 1). Sites were located at similar distance from shore (between 5.8 and 3.8 km), with available habitat dominated by hard-coral communities (ca. 40-55% live coral cover, Burt et al. 2011a) and were at 6 – 7m in depth (Grizzle et al. 2015).

**Fig. 1.**
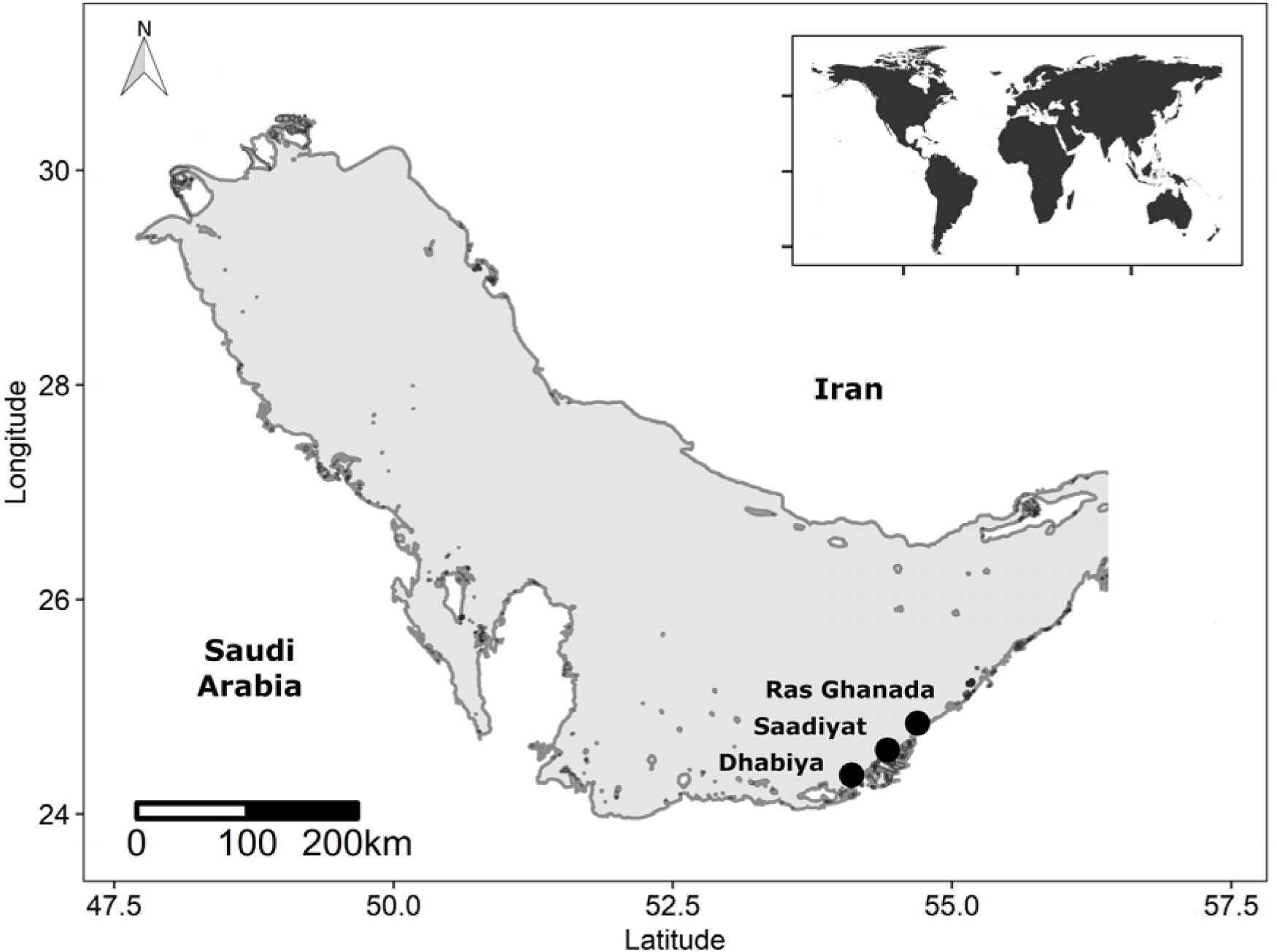
Study area and sampling locations in the southern part of the Persian Gulf. DHAB: Dhabiya; SAAD: Saadiyat; RASG: Ras Ghanada.

#### In situ *behavioural observation*

Observations were undertaken by scuba diving on cloudless days between 10:00 and 14:00, reducing any influence of changing light levels on behaviour (Layton and Fulton 2014). In each site, across seasons, replicate coral heads holding *P. trichrourus* individuals were identified by visual searches made whilst swimming along haphazardly located transects of the reef. Coral heads held one (56% of coral heads examined), two (34%), three (7%) or four (2%) adult *P. trichrourus*. An individual *P. trichrourus* was haphazardly selected and its territory identified and marked using numbered flags (Magalhaes et al. 2013). When several adults shared the same coral head, in order to minimize the developmental variability among individuals being sampled, the individual with the largest body size was selected. Following marking of the territory, focal fish were allowed to acclimate for 4 min (), and then filmed for 2 minutes (using DBPOWER HD 1080P). A minimum distance of 3 m was maintained between observer and subject. Observations were aborted if fish displayed any adverse reactions to diver’s presence (< 5% of individuals). Across sites and seasons, a total of 226 individual behavioural observations were recorded (winter: Dhabiya n = 26, Saadiyat n = 32, Ras Ghanada n = 25; spring: Dhabiya n = 26, Saadiyat n = 32, Ras Ghanada n = 25; summer: Dhabiya n = 23, Saadiyat n = 31, Ras Ghanada n = 29).

#### *Analyses of video quantifying* in situ *individual behaviour*

Mean distance from refugia was defined as the distance from the ‘home’ coral every ten seconds (Beck et al. 2016) and estimated to the nearest 5 cm referenced against the territory marking flag’s length (25 cm) and the individual’s body size (taken as 5 cm SL). Resting was taken as total time (seconds) spent immobile (i.e. not active swimming, foraging or chasing behaviour) across each 2-minute video. Feeding rate was the total number of bites during the observation period, while diet was identified by classifying each food bite as either on pelagic or benthic substrates. Benthic food bites were further classified as being on live coral, coral rubble, turf or sand. Behavioural data were initially extracted from 15% of videos and independently analysed and validated by two observers. As results differed by < 5 % between observers, all remaining video analyses were carried out by one observer (DD).

#### In situ *environmental measurements*

To assess seasonal changes in abiotic variables in the Gulf, across the dates of all behavioural observations, daily *in situ* SST (°C) and dissolved oxygen (DO, mg l^-1^) measurements were taken at 1200 hrs at each site using a YSI Professional Plus multi-parameter probe, while underwater visibility (in m) was visually estimated *in situ*. Chlorophyll-a (chl-a, mg l^-1^) measurements were downloaded as monthly composite data at 0.05 resolution from the MODIS-aqua satellite (NASA, Ocean Ecology Laboratory), while salinity (psu), wind (m s^-1^) and currents (m s^-1^) were obtained from the NCEP climate forecast system version 2 (CFSv2, Saha et al 2011). Salinity and current data were accessible as daily means (resolution 0.25- deg × 0.25-deg), while wind data were taken at 1200 hrs (0.205-deg × ∼0.204-deg) across all experimental days.

To estimate the relative frequency of predation or competition across seasons and sites, the abundance of predators (i.e. piscivorous fish with SL ≥ 15 cm observed interacting with *P. trichrourus*) and competitors (i.e. fishes with comparable ecology/body size to *P. trichrourus* observed chasing/being chased by *P. trichrourus*) entering each individual territory was quantified for each observation.

### Aquarium-based behavioural quantification

To experimentally determine the impact of water temperature on behaviour, we examined time away from refugia and distance moved of *P. trichrourus* in controlled conditions in the laboratory. Sixteen adults (mean body size 4.78 [± 0.13 SE] cm SL) were collected from Saadiyat during winter. Eight individuals were randomly allocated to separate pierced 6L bottles within either of two 900L holding tanks (control and experimental group). Each bottle had a 10-cm-long plastic pipe provided for shelter. All tanks were kept at 21°C, 40 psu, 8.15 pH, 6.5 mg l^-1^ DO and 12 hours dark/light cycle reflecting *in situ* winter condition. 25% water changes were performed weekly, with ammonia, nitrite and nitrate kept at 0 ppt. Fish were fed *ad libitum* twice daily with commercial pellets.

After a week for acclimation, refuge use and movement of all individuals was tested at 21°C. To do this a randomly chosen individual had its holding container submerged in an experimental tank (80 L * 50 W *40 H [cm], filled to 20 cm H) and was left to acclimate for 10 minutes. The tank’s base was divided into 40 squares (10 * 10 cm grid), a 10-cm-long segment of plastic pipe was provided as shelter, and a DBPOWER HD 1080P video camera was placed one metre above the tank. Following acclimation the individual was released and led to the shelter; a transparent bottomless bottle was used to confine the individual which was then left for 20 minutes. Subsequently the bottle was removed and behaviour videoed for 16 minutes. All videoing occurred between 1000 hrs and 1400 hrs, with a 25% water change between individuals. Water temperature within the experimental holding tank was increased to 28°C (∼0.5°C/6 h) across four days, while the temperature within the control remained constant (21 °C). Fish were then left for 7 days. As described above, all individuals were then individually videoed and behaviours experimentally examined.

Within experimental videos, the first 5 minutes were discarded to allow time of acclimation after activation of the video camera. Time away from refuge was quantified (in seconds) when individuals moved 10 cm from the shelter. Distance moved was quantified (in cm) by counting the number of grid-lines crossed over 11 min with one cross being assumed to equal to 10 cm movement, independently of the distance from refuge.

### Statistical Analysis

To determine whether abiotic and/or biotic factors significantly differed among seasons, Kruskal-Wallis test was undertaken. Abiotic factors were then combined using Principal Components Analysis, to identify the main axes of environmental variation among observations; axes that explained more than 15% of the total variance (PC1 to PC3, hereafter ‘environmental PCs’) were retained and used in further analyses (Magalhaes et al. 2016).

To determine the impact of season, site, environmental PCs and predator/competitor abundance on distance from refugia and resting time, a Generalised Linear Mixed Model (GLMM) with Gaussian errors was used, while feeding rate against all predictor variables (as above) was examined using a GLMM with Poisson error distribution and log link function. Both distance from refugia and resting were not normally distributed and were, respectively, log and log (X + 1) transformed prior to testing. In all tests, date of sampling was fitted as a random effect and backwards model selection followed, using likelihood ratio tests to examine the significance of each term removed from the model. To determine whether food preference significantly differed between seasons, individual bites on substrate were analysed using Kruskal-Wallis test.

To determine the effect of experimental change in temperature on time away from refugia and distance moved, behavioural data before and after temperature manipulation from the control and experimental groups were compared using Mann-Whitney U tests. All data were analysed in R (R core development team 3.5.1, 2018).

## Results

### *In situ* environmental variation

Significant differences among seasons in SST, DO, chl-a, salinity and visibility were apparent (Kruskal-Wallis test: p < 0.05 in all cases; Table S1). SST was highest in summer and lowest in winter, while DO showed the opposite trend. Chl-a concentration was higher in summer than the other seasons. PCA showed that 82% of the variation in the raw abiotic variables was captured by the first three principal components, with a high score in PC1 (39%) being associated with low SST and chl-a, and high DO. A high score in PC2 (25%) was associated with high salinity and low visibility, and a high score in PC3 (18%) was associated with high water current (Table 1).

**Table 1.** Loadings of each environmental variables on the first three PC axes and the percent variance explained by each axis. The highest loadings on each of the first three PCs are showed in bold.

| Factors | PC1 | PC2 | PC3 |
| --- | --- | --- | --- |
| Temperature | **-0.58** | -0.06 | -0.04 |
| Dissolved Oxygen | **0.58** | 0.11 | 0.06 |
| Chl-a | **-0.49** | 0.25 | 0.03 |
| Salinity | -0.10 | **0.70** | 0.14 |
| Wind speed | -0.25 | -0.22 | -0.26 |
| Water current | 0.06 | -0.14 | **-0.83** |
| Visibility | -0.08 | **-0.60** | **0.47** |

Across all sites and seasons, predator abundance was dominated by *Lutjanus ehrenbergii*, comprising 97% of counts (Table S2). Predator abundance significantly differed among seasons (Kruskal-Wallis test, H = 73.790, df = 2, p < 0.001), with predators being more abundant in summer than spring or winter, while among sites predators were more abundant in Dhabiya than Saadiyat or Ras Ghanada (Fig. S1). The predominant competitors were other *P. trichrourus* and *Pomacentrus aquilis* individuals. Competitor abundance significantly differing among seasons (H = 7.191, df = 2, p = 0.027), with slightly higher abundance in spring than winter (Table S2).

### *In situ* behavioural analysis

Season had a significant effect on *P. trichrourus* distance from refugia (Table 2), with individuals showing a higher distance from refugia in spring than in either winter or summer (Table S3). Although site itself did not have a significant effect on distance from refugia, there was a significant interaction between site and season (Table 2), with individuals in Ras Ghanada and Saadiyat showing higher distance from refugia in spring than winter or summer compared to individuals from Dhabiya (Fig. 2a). Overall, *P. trichrourus* spent more time resting during summer and winter than spring (Table S3). However, only the interaction between site and season had a significant effect on resting time (Table 2); a clear pattern in seasonal differences in resting was evident only in Ras Ghanada and Saadiyat compared to Dhabiya (Fig. 2b). Season and the interaction between site and season had a significant effect on individuals’ feeding behaviour (Table 2), with bite rate being twice as high in spring as in summer, and four times higher in spring than winter (Table S3). Clear seasonal changes in feeding rate were apparent in Ras Ghanada and Saadiyat, while in Dhabiya the differences between spring and summer were less clear-cut (Fig. 2c).

**Table 2.** Results of GLMM analysis of effects of biotic/abiotic factors on *Pomacentrus trichrourus’ in situ* behaviour. Gaussian error structure for distance from refugia and resting time; Poisson error distribution and with log link function for feeding rate. Significant p values are highlighted in bold.

| Response | Predictor | Residual deviance<sub>(df)</sub> | Change in deviance<sub>(df)</sub> | p |
| --- | --- | --- | --- | --- |
| **Distance from refugia** | Season | 8.298<sub>(5)</sub> | 10.925<sub>(2)</sub> | **0.004** |
|  | Site | -1.1050<sub>(5)</sub> | 1.522<sub>(2)</sub> | 0.467 |
|  | Site : season | -2.627<sub>(7)</sub> | 30.565<sub>(4)</sub> | **< 0.001** |
|  | PC1 | -37.767<sub>(14)</sub> | 0.430<sub>(1)</sub> | 0.512 |
|  | PC2 | -33.192<sub>(11)</sub> | 3.690<sub>(1)</sub> | 0.055 |
|  | PC3 | -37.390<sub>(13)</sub> | 0.377<sub>(1)</sub> | 0.539 |
|  | Predators | -36.882<sub>(12)</sub> | 0.508<sub>(1)</sub> | 0.476 |
|  | Competitors | -38.197<sub>(15)</sub> | 0.176<sub>(1)</sub> | 0.674 |
| **Resting time** | Season | 89.580<sub>(5)</sub> | 4.384<sub>(2)</sub> | 0.112 |
|  | Site | 86.431<sub>(5)</sub> | 0.539<sub>(6)</sub> | 0.539 |
|  | Site : season | 85.196<sub>(7)</sub> | 29.556<sub>(4)</sub> | **< 0.001** |
|  | PC1 | 54.937<sub>(12)</sub> | 1.013<sub>(1)</sub> | 0.314 |
|  | PC2 | 55.640<sub>(11)</sub> | 0.703<sub>(1)</sub> | 0.408 |
|  | PC3 | 53.924<sub>(13)</sub> | 1.324<sub>(1)</sub> | 0.25 |
|  | Predators | 52.042<sub>(15)</sub> | 0.015<sub>(1)</sub> | 0.908 |
|  | Competitors | 52.601<sub>(14)</sub> | 0.558<sub>(1)</sub> | 0.455 |
| **Feeding rate** | Season | 1730.2<sub>(4)</sub> | 31.427<sub>(2)</sub> | **< 0.001** |
|  | Site | 1700.7<sub>(4)</sub> | 1.883<sub>(2)</sub> | 0.39 |
|  | Site : season | 1698.8<sub>(6)</sub> | 17.959<sub>(4)</sub> | **0.001** |
|  | PC1 | 1674.6<sub>(12)</sub> | 1.459<sub>(1)</sub> | 0.227 |
|  | PC2 | 1680.8<sub>(10)</sub> | 3.805<sub>(1)</sub> | 0.051 |
|  | PC3 | 1677.0<sub>(11)</sub> | 2.487<sub>(1)</sub> | 0.115 |
|  | Predators | 1672.9<sub>(14)</sub> | 0.137<sub>(1)</sub> | 0.137 |
|  | Competitors | 1673.1<sub>(13)</sub> | 0.165<sub>(1)</sub> | 0.685 |

**Fig. 2.**
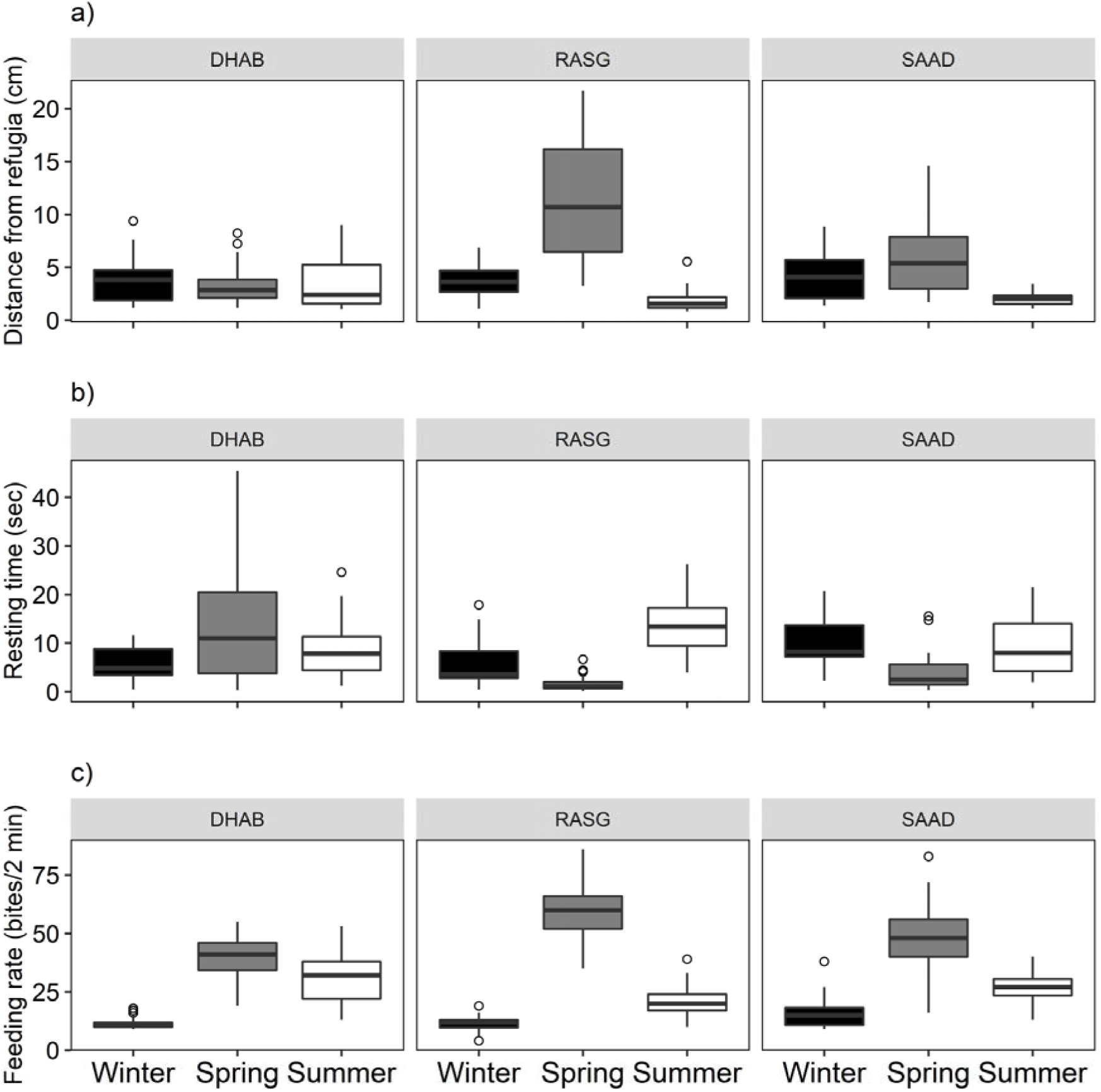
Changes in *Pomacentrus trichrourus* behaviour *in situ* among seasons (winter = black, spring = grey, summer = white) and locations (Dhabiya [DHAB], Ras Ghanada [RASG] and Saadiyat [SAAD]): **(a)** mean distance from refugia (cm); **(b)** time spent resting (s/120 s) and **(c)** number of feeding bites in 120 s. Thick horizontal lines show the median, boxes show inter-quartile (IQR). Whiskers indicate the range of data, and dots show outliers that are more than 1.5 IQR above the 75^th^ percentile, or more than 1.5 IQR below the 25^th^ percentile.

Significant differences in diet were apparent among seasons, with planktonic prey representing 93% and 96% of food bites in winter and spring, respectively, while only accounting for 73% in summer (Kruskal-Wallis, H =139.04, df = 2, P < 0.001). Benthic food bites were predominantly given on live corals (5 and 4% of total food bites in winter and spring, 17% in summer) while coral rubble, turf and sand accounted for less than 1% of total food bites in winter and spring, and ≤4% in summer.

### Laboratory behavioural assays

Before temperature manipulation there were no differences between the control and the experimental group in both time away from refugia (Mann-Whitney U test, W = 44, p = 0.14) and distance moved (W = 40, p = 0.41); however, differences in time away from refugia and distance moved between the control and the experimental group become significant post temperature manipulation (W = 2, p= 0.001 and W = 3, p = 0.002, respectively) (Fig. 3a, b).

**Fig. 3.**
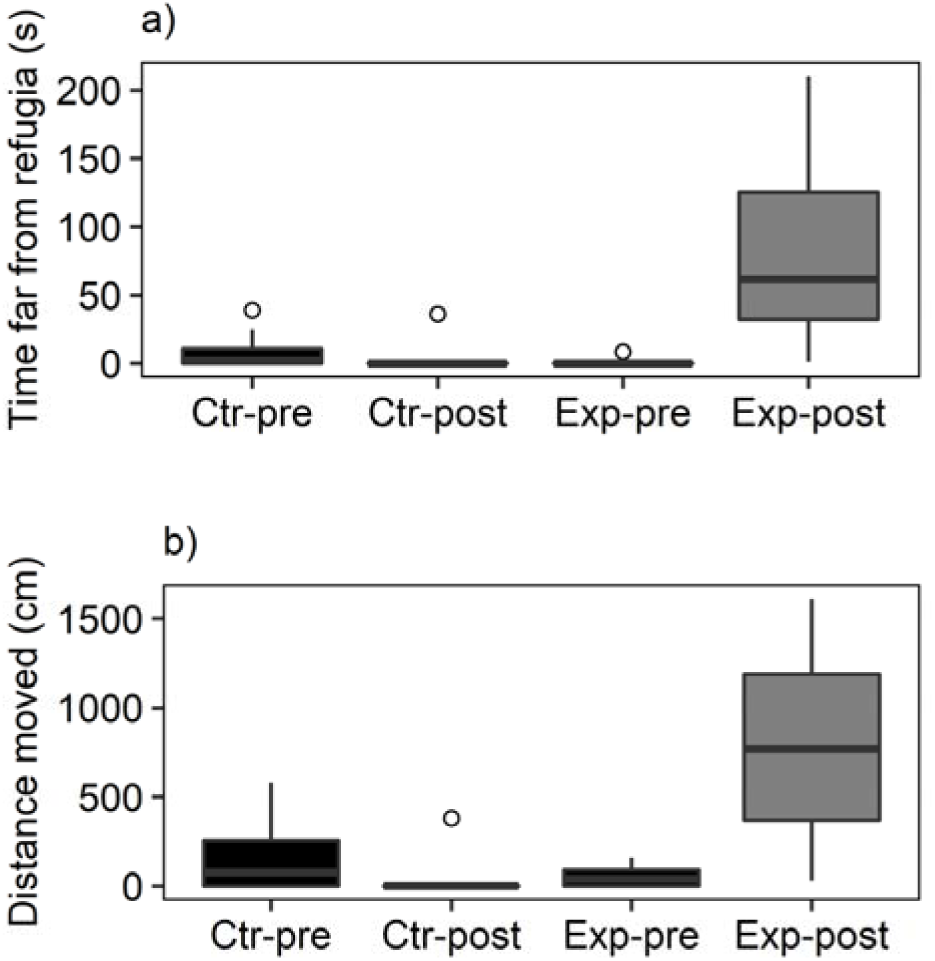
Changes in *Pomacentrus trichrourus* behaviour ex *situ* among temperature treatment: mean time spent far from refugia (s), and **(b)** total distance moved (cm). Colours indicate temperature (black = 21°C, grey = 28°C). Thick horizontal lines show the median, boxes show inter-quartile (IQR). Whiskers indicate the range of data, and dots show outliers that are more than 1.5 IQR above the 75^th^ percentile, or more than 1.5 IQR below the 25^th^ percentile.

## Discussion

Although increasing variance in global SSTs may directly impact coral reef fish community biodiversity (Pratchett et al. 2015; Pecl et al. 2017; Hughes et al. 2018), coral reef fish populations might be able to mitigate the effect of rapidly changing environments through behavioural plasticity (Nagelkerken and Munday 2016; Scott et al. 2017; Shraim et al. 2017; Chase et al. 2018). We examined the impact of seasonal changes in biotic and abiotic conditions on distance from refugia, activity and feeding ecology of the pale-tail damselfish *P. trichrourus* within the southern Gulf. We found that *P. trichrourus* substantially increased resting time and reduced distance from refugia and feeding rate at sub-optimal environmental temperatures (both high and low) when observed *in situ* and when tested in the laboratory. Individuals were also observed to shift from a planktonic-based diet to a mixed planktonic and benthic diet during the Gulf’s extreme summer season. Our results show considerable scope for behavioural and trophic plasticity in *P. trichrourus*, which may allow this species to cope with seasonal variance in environmental conditions, including those that are above the mortality thresholds (Nilsson et al. 2009; Munday et al. 2009; Rummer et al. 2014; Rodgers et al. 2018) and below the optimum temperature of Indo-Pacific reef fish (Eme and Bennett 2008; Figueira and Booth 2010; Nakamura et al. 2013).

We observed substantial behavioural and trophic plasticity in *P. trichrourus’* populations among seasons. In particular, *P. trichrourus* were observed to reduce distance from refugia and feeding rate and increase resting time in winter and summer compared to spring. These seasonal changes are plausibly related to variation in water temperature, as changes in environmental temperature are known to have a direct effect on fish physiology (Angilletta 2009; Abram 2017). In ectothermic animals, environmental temperatures departing from the optimum are associated with reductions in aerobic scope, reducing energy for costly activities (i.e. swimming, feeding, chasing, etc.) (Pörtner and Knust, 2007; Pörtner and Farrell 2008; Munday 2012). The reduction in activity observed in *P. trichrourus,* is in line with recent work suggesting that, when exposed to short-term higher than average SSTs, coral reef fish may temporarily mitigate bioenergetic inefficiency through reduction in energetically costly activities, as well as through changes in diet (Nowicki et al. 2012; Johansen et al. 2014; Scott et al. 2017; Chase et al. 2018). However, such strategies may not be sufficient to sustain prolonged periods (i.e. months) of extreme temperature (Rummer et al. 2014; Rodgers et al. 2018). For example, reductions in food intake and quality, coupled with low aerobic scope under extreme temperatures, can disrupt an ectothermic organism’s energy balance, with substantially less energy available for long-term maintenance, growth and reproduction (Gillooly et al. 2001; Pörtner et al. 2010; Neuheimer et al. 2011), impacting individual fitness and potentially survival (Chase et al. 2018).

*Pomacentrus trichrourus’s* persistence within the Gulf implies that they have adapted over the long-term to the extreme temperature conditions experienced there (Rummer and Munday 2017). Our results hint that the key to their survival may be their ability to undertake seasonal energy recovery in spring, compensating for energy loss during winter while building up energetic stores to endure the upcoming summer season (Armstrong and Bond 2013). Many studies on temperate fishes have highlighted the ability to compensate energetic and growth losses during periods of adverse temperatures or food scarcity through increased feeding rate and feeding activity when conditions become favourable again (Sevgili et al. 2013; Armstrong and Bond 2013; Furey et al. 2016; Peng et al. 2017). For *P. trichrourus* such periods of recovery, where it can access highly nutritious food resources (Shraim et al. 2017) under optimal environmental temperatures (Kaschner et al 2016), may be vital for mitigating energy loss during winter and summer, and hence ultimately permit the persistence of abundant populations within the southern Gulf throughout the year.

Our lab experiments are consistent with the idea that the seasonal differences in behaviour observed in the field were the result of changes in temperature. However, other potentially important environmental variables also changed with the seasons, including predators and primary productivity. These variables could plausibly be directly responsible for changes in fish behaviour. For example, predators might have had a negative effect on *P. trichrourus* movement (i.e. reduced distance from refugia) and feeding rate (Dill & Fraser 1984; Beck et al. 2016; Catano et al. 2016), potentially leading to the shift to mixed planktonic and benthic diet observed in summer, when *P. trichrourus* may have to rely more on safer benthic food resources (i.e. live corals), despite the loss of nutritional value (Shraim et al. 2017). On the contrary, we did not observe any correlation between changes in primary productivity and changes in feeding activity, suggesting that shifts in diet may not be related to primary productivity (measured as chlorophyll-a concentration) (Rueda et al. 2015; Zhou et al. 2016) but the result of temperature-related differences in food preference (Shraim et al. 2017). Ultimately, we would need manipulative experiments to confirm the potential for predators or other drivers to produce similar behavioural changes to those seen in our temperature manipulation.

The Gulf’s average SST is increasing twice as fast as the global average (Al-Rashidi et al. 2009), with predictions of further increases of 0.5°C – 1.4°C by 2050, associated with hot salt brine discharge derived from local desalinisation plants (AGEDI 2016; Vaughan et al. 2019). As *P. trichrourus* populations live for five months of the year above their predicted thermal optimum (Jun–Oct, mean SST >30°C) (Kaschner et al. 2016), of which two months encompass temperatures above the mortality threshold of low latitude Indo-Pacific damselfish (July and August, mean SST ≥ 33°C) (Nilsson et al. 2009; Munday et al. 2009; Rummer et al. 2014; Rodgers et al., 2018), Gulf populations may already be living close to their upper thermal limits; further increases in temperature are expected to pose a serious threat to population persistence. In addition, *P. trichrourus* populations in the Gulf are already considered endangered, associated with a fragmented distribution and strong dependence to diminishing coral habitat resources (Buchanan et al. 2016; 2019). Indeed, *Pomacentrus trichrourus* are highly associated with live coral colonies, utilising them for recruitment, shelter and as trophic resources (Buchanan et al. 2016; Shraim et al. 2017). Therefore, any increases in SST that pose a serious threat to local coral communities, where bleaching events and consequent coral mortality are already common (Riegl and Purkis 2015; Riegl et al. 2018), may deprive *P. trichrourus* populations of suitable habitat and food resources (Keith et al. 2018; Pratchett et al. 2018).

The Gulf’s marine community encompasses 241 species of coral-associated bony fish, which are considered a biogeographic subset of the Indian Ocean’s fauna that re-colonised the region between 6000 and 9000 years ago (Riegl and Purkis 2012). The coral reef fish community richness in the Gulf is thought to be lower than that in the Indian Ocean as a consequence of the Gulf’s physical extremes, which fewer Indian Ocean species are able to tolerate (Coles 2003; Feary et al. 2010). There is still little understanding of how tolerance is achieved in Gulf-dwelling species. For *P. trichrourus* this study has shown that populations may be able to behaviourally mitigate the high variance in temperature and temperature extremes which are consistent with predictions for the tropical ocean by the end of the century (ICCP 2014). It appears that by reducing movement from shelter and feeding activities within extreme seasons (summer, winter) and showing metabolic compensatory activity behaviour (including feeding) during the environmentally benign spring season, populations of *P. trichrourus* are able to thrive in one of the most thermally extreme marine environments. Such adaptive behavioural plasticity is consistent with current theory regarding how animals deal with extremes, and may be an important mechanism for how increases in temperature and temperature variability may be mitigated by fishes on Indo-Pacific reefs by the end of the century.

## Acknowledgments

All research was carried out under approval of the NYU – AD Institutional Animal Care and Use Committee (IACUC protocol No. 16-0005) and according to the University’s animal ethics guidelines. Fish collection around Abu Dhabi’s reefs were carried with permission of the Environment Agency, Abu Dhabi (protocol No. EAD-TMBS-RP-0). Thanks to the University of Nottingham for support thought the whole study. Thanks to Dain McParland and Noura Al-Mansoori for field and lab support.

On behalf of all authors, the corresponding author states that there is no conflict of interest.

## Electronic Supplementary Material

**Table S1.** Differences in abiotic (a) and biotic (b) environmental variables between seasons.

| Environmental factor | Winter | Spring | Summer | Kruskal-Wallis |
| --- | --- | --- | --- | --- |
| a) *Abiotic* | n= 4 | n = 6 | n = 6 |  |
| **SST** | 20.60 (0.12) | 27.45 (0.19) | 33.85<br>(0.11) | H = 13.294, df = 2, p = **0.001** |
| **DO** | 6.54 (0.04) | 5.61 (0.02) | 4.86 (0.01) | H = 13.274, df = 2, p = **0.001** |
| **Chl-a** | 1.38 (0.05) | 1.38 (0.17) | 2.97 (0.62) | H = 10.638, df = 2 , p = **0.005** |
| **Salinity** | 39.93 (0.05) | 38.93 (0.03) | 39.85<br>(0.13) | H = 10.770, df = 2, p = **0.005** |
| **Wind speed** | 5.5 (0.70) | 5.91 (0.34) | 5.57 (0.56) | H = 2.592, df = 2, p = 0.274 |
| **Water current** | 0.03 (0.02) | 0.03 (0.01) | 0.03 (0.01) | H = 1.456, df = 2 , p = 0.483 |
| **Visibility** | 6.00 (1.11) | 11.95 (1.23) | 8.25 (0.66) | H = 6.231, df = 2, p = **0.044** |
| b) *Biotic* | n = 61 | n = 83 | n = 83 |  |
| **Predators** | 0.22 (0.1) | 0.92 (0.2) | 6.41 (1) | H = 73.790, df = 2, p < **0.001** |
| **Competitors** | 0.38 (0.1) | 0.75 (0.1) | 0.49 (0.1) | H = 7.191, df = 2, p = **0.027** |

**Table S2.** Comparison of mean (± SE), maximum and total of predators (**a**) and competitors that the focal species *Pomacentrus trichrourus* encountered during the 2 minutes behavioural observation in winter (left), spring (middle) and summer (right). Species ranked by order of abundance at the summer season.

| Species | Winter |  |  | Spring |  |  | Summer |  |  |
| --- | --- | --- | --- | --- | --- | --- | --- | --- | --- |
|  | Mean | Max | n | Mean | Max | n | Mean | Max | n |
| **(a) Predators** |  |  |  |  |  |  |  |  |  |
| *Lutjanus ehrenbergii* | 0.08<br>(0.05) | 3 | 5 | 0.89<br>(0.21) | 10 | 74 | 6.12<br>(1.00) | 59 | 508 |
| *Scolopsis ghanam* | 0.12<br>(0.06) | 3 | 7 | 0.01<br>(0.01) | 1 | 1 | 0.17<br>(0.06) | 3 | 14 |
| *Epinephelus spp* | 0 | 0 | 0 | 0.02<br>(0.02) | 1 | 1 | 0.09<br>(0.05) | 1 | 6 |
| *Lutjanus fulviflamma* | 0 | 0 | 0 | 0.02<br>(0.02) | 1 | 1 | 0.09<br>(0.05) | 1 | 3 |
| **Total** | **0.22**<br>**(0.10)** |  | **12** | **0.92**<br>**(0.20)** |  | **76** | **6.41**<br>**(1.00)** |  | **525** |
| **(b) Competitors** |  |  |  |  |  |  |  |  |  |
| *Pomacentrus trichrourus* | 0.35<br>(0.07) | 2 | 21 | 0.75<br>(0.01) | 4 | 62 | 0.45<br>(0.06) | 2 | 37 |
| *Pomacentrus aquilis (j)* | 0.05<br>(0.04) | 2 | 3 | 0.59<br>(0.07) | 3 | 49 | 0.3<br>(0.05) | 2 | 25 |
| *Pomacentrus aquilis* | 0.03<br>(0.02) | 1 | 2 | 0 | 0 | 0 | 0.05<br>(0.02) | 1 | 4 |
| **Total** | **0.38**<br>**(0.10)** |  | **26** | **0.75**<br>**(0.10)** |  | **111** | **0.49**<br>**(0.10)** |  | **66** |

**Table S3.** Comparison of *in situ* behaviour (median ± SE, max and min) between seasons.

| Behaviour | Winter (n = 60) |  |  | Spring (n = 83) |  |  | Summer (n = 83) |  |  |
| --- | --- | --- | --- | --- | --- | --- | --- | --- | --- |
|  | Median | Max | Min | Median | Max | Min | Median | Max | Min |
| **Distance from refugia** | 3.77<br>(0.3) | 9.38 | 1.0<br>8 | 5.38<br>(0.5) | 21.6<br>9 | 1.1<br>5 | 1.96<br>(0.2) | 9 | 0.85 |
| **Resting** | 5.62<br>(0.6) | 20.7<br>4 | 0.4<br>8 | 2.48<br>(1.0) | 45.4<br>5 | 0.1<br>2 | 10.37<br>(0.7) | 26.2<br>4 | 10.3<br>7 |
| **Feeding rate** | 12<br>(0.7) | 38 | 4 | 49<br>(0.9) | 86 | 16 | 25 (0.9) | 53 | 25 |

**Fig. S1.**
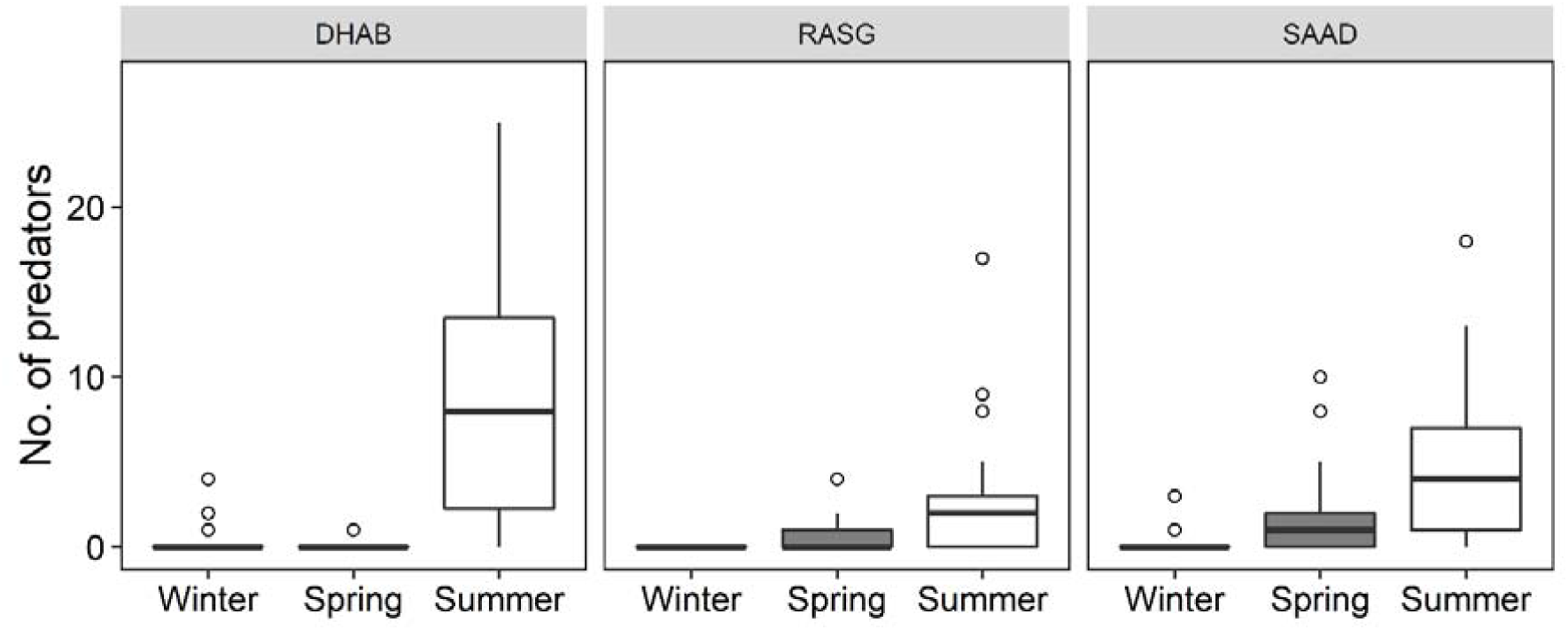
Changes in abundance of *Pomacentrus trichrourus’* potential predators with seasons (winter = black, spring = grey, summer = white) and locations (Dhabiya [DHAB], Ras Ghanada [RASG] and Saadiyat [SAAD]). Thick horizontal lines show the median, boxes show inter-quartile (IQR). Whiskers indicate the range of data, and dots show outliers that are more than 1.5 IQR above the 75^th^ percentile, or more than 1.5 IQR below the 25^th^ percentile.

## Reference List

Abram PK, Boivin G, Moiroux J, Brodeur J (2017) Behavioural effects of temperature on ectothermic animals: unifying thermal physiology and behavioural plasticity. Biol Rev 92:1859–1876

Allen GR (1991) Damselfishes of the world. Mergus Publishers, Melle, Germany, p 271

AGEDI (2016) Final technical report: Regional desalination and climate change (report: CCRG/IO). Abu Dhabi: Local, National, and Regional Climate Change Programme, Abu Dhabi Global Environmental Data Initiative (AGEDI).

Al-Rashidi TB, El-Gamily HI, Amos CL, Rakha KA (2009) Sea surface temperature trends in Kuwait Bay, Arabian Gulf. Nat Hazards 50:73–82

Albouy C, Leprieur F, Le Loc’h F, Mouquet N, Meynard CN, Douzery EJP, Mouillot D (2015) Projected impacts of climate warming on the functional and phylogenetic components of coastal Mediterranean fish biodiversity. Ecography (Cop) 38:681–689

Angilletta MJ (2009) Thermal Adaptation: A Theoretical and Empirical Synthesis. Oxford University Press, New York.

Armstrong JB, Bond MH (2013) Phenotype flexibility in wild fish: Dolly Varden regulate assimilative capacity to capitalize on annual pulsed subsidies. J Anim Ecol 82:966–975

Beck HJ, Feary DA, Fowler AM, Madin EMP, Booth DJ (2016) Temperate predators and seasonal water temperatures impact feeding of a range expanding tropical fish. Mar Biol 163:1–14

Bellard C, Bertelsmeier C, Leadley P, Thuiller W, Courchamp F (2012) Impacts of climate change on the future of biodiversity. Ecol Lett 15:365–377

Buchanan JR, Krupp F, Burt JA, Feary DA, Ralph GM, Carpenter KE (2016) Living on the edge: Vulnerability of coral-dependent fishes in the Gulf. Mar Pollut Bull 105:480–488

Buchanan JR, Ralph GM, Krupp F, Harwell H, Abdallah M, Abdulqader E, Al-Husaini M, Bishop JM, Burt JA, Choat JH, Collette BB, Feary DA, Hartmann SA, Iwatsuki Y, Kaymaram F, Larson HK, Matsuura K, Motomura H, Munroe T, Russell B, Smith-Vaniz W, Williams J, Carpenter KE (2019) Regional extinction risks for marine bony fishes occurring in the Persian/Arabian Gulf. Biol Conserv 230:10–19

Burt JA, Al-Harthi S, Al-Cibahy A (2011a) Long-term impacts of coral bleaching events on the world’s warmest reefs. Mar Environ Res 72:225–229

Burt JA, Feary DA, Bauman AG, Usseglio P, Cavalcante GH, Sale PF (2011b) Biogeographic patterns of reef fish community structure in the northeastern Arabian Peninsula. ICES J Mar Sci 68:1875–1883

Candolin U (2018) Adaptedness of Behavior. In: Vonk J, Shackelford T (eds) Encyclopedia of Animal Cognition and Behavior. Springer, Cham, pp 1–11

Catano LB, Rojas MC, Malossi RJ, Peters JR, Heithaus MR, Fourqurean JW, Burkepile DE (2016) Reefscapes of fear: Predation risk and reef hetero-geneity interact to shape herbivore foraging behaviour. J Anim Ecol 85:146–156

Chase TJ, Nowicki JP, Coker DJ (2018) Diurnal foraging of a wild coral-reef fish *Parapercis australis* in relation to late-summer temperatures. J Fish Biol 153–158

Cheal AJ, MacNeil MA, Emslie MJ, Sweatman H (2017) The threat to coral reefs from more intense cyclones under climate change. Glob Chang Biol 23:1511–1524

Coles SL, Fadlallah YH (1991) Reef coral survival and mortality at low temperatures in the Arabian Gulf: new species-specific lower temperature limits. Coral Reefs 9:231–237

Coles SL (2003) Coral species diversity and environmental factors in the Arabian Gulf and the Gulf of Oman: A comparison to the Indo-Pacific region. Atoll Res Bull 1–19

Côté IM, Darling ES (2010) Rethinking ecosystem resilience in the face of climate change. PLOS Biol 8: e1000438

Day PB, Stuart-Smith RD, Edgar GJ, Bates AE (2018) Species’ thermal ranges predict changes in reef fish community structure during 8 years of extreme temperature variation. Divers Distrib 24:1036–1046

Dill LM, Fraser AHG (1984) Risk of predation and the feeding behavior of juvenile coho salmon (*Oncorhynchus kisutch*). Behav Ecol and Sociobiol 16:65–71

Doney SC, Ruckelshaus M, Emmett Duffy J, Barry JP, Chan F, English CA, Galindo HM, Grebmeier JM, Hollowed AB, Knowlton N, Polovina J, Rabalais NN, Sydeman WJ, Talley LD (2012) Climate Change Impacts on Marine Ecosystems. Ann Rev Mar Sci 4:11–37

Eme J, Bennett WA (2008) Low temperature as a limiting factor for introduction and distribution of Indo-Pacific damselfishes in the eastern United States. J Therm Biol 33:62–66

Feary DA, Burt JA, Bauman AG, Usseglio P, Sale PF, Cavalcante GH (2010) Fish communities on the world’s warmest reefs: What can they tell us about the effects of climate change in the future? J Fish Biol 77:1931–1947

Figueira WF, Biro P, Booth DJ, Valenzuela VC (2009) Performance of tropical fish recruiting to temperate habitats: Role of ambient temperature and implications of climate change. Mar Ecol Prog Ser 384:231–239

Figueira WF, Booth DJ (2010) Increasing ocean temperatures allow tropical fishes to survive overwinter in temperate waters. Glob Chang Biol 16:506–516

Folguera G, Bastías DA, Caers J, Rojas JM, Piulachs MD, Bellés X, Bozinovic F (2011) An experimental test of the role of environmental temperature variability on ectotherm molecular, physiological and life-history traits: Implications for global warming. Comp Biochem Physiol - A Mol Integr Physiol 159:242–246

Furey NB, Hinch SG, Mesa MG, Beauchamp DA (2016) Piscivorous fish exhibit temperature-influenced binge feeding during an annual prey pulse. J Anim Ecol 85:1307–1317

Gillooly JF, Brown JH, West GB, Savage VM, Charnov EL (2001) Effects of size and temperature on metabolic rate. Science 293:2248–2251

Gordon TAC, Harding HR, Clever FK, Davidson IK, Windsor FM, Armstrong JD, Bardonnet A, Bergman E, Britton JR, Côté IM, D’agostino D, Greenberg LA, Harborne AR, Kahilainen KK, Metcalfe NB, Mills SC, Milner NJ, Mittermayer FH, Montorio L, Nedelec SL, Prokkola JM, Rutterford LA, Salvanes AG, Simpson SD, Vainikka A, Pinnegar JK, Santos EM (2018) Fishes in a changing world: learning from the past to promote sustainability of fish populations. J Fish Biol 92:804–827

Grizzle RE, Ward KM, AlShihi RMS, Burt JA (2015) Current status of coral reefs in the United Arab Emirates: Distribution, extent, and community structure with implications for management. Mar Pollut Bull 105:515–523

Hughes TP, Kerry JT, Baird AH, Connolly SR, Dietzel A, Eakin CM, Heron SF, Hoey AS, Hoogenboom MO, Liu G, McWilliam MJ, Pears RJ, Pratchett MS, Skirving WJ, Stella JS, Torda G (2018) Global warming transforms coral reef assemblages. Nature 556:492– 496

IPCC (2012) Managing the risks of extreme events and disasters to advance climate change adaptation. In: A Special Report of Working Groups I and II of the Intergovernmental Panel on Climate Change (eds Field CB, Barros V, Stocker TF et al.), Cambridge University Press, Cambridge, NY, USA.

IPCC (2014) Summary for Policy Makers. Clim Chang 2014 Impacts, Adapt Vulnerability - Contrib Work Gr II to Fifth Assess Rep 1–32

Johansen JL, Messmer V, Coker DJ, Hoey AS, Pratchett MS (2014) Increasing ocean temperatures reduce activity patterns of a large commercially important coral reef fish. Glob Chang Biol 20:1067–1074

Kaschner K, Kesner-Reyes K, Garilao C, Rius-Barile J, Rees T, Froese R (2016). AquaMaps: predicted range maps for aquatic species. World wide web electronic publication, www.aquamaps.org, Version 08/2016

Keith SA, Baird AH, Hobbs J-PA, Woolsey ES, Hoey AS, Fadli N, Sanders NJ (2018) Synchronous behavioural shifts in reef fishes linked to mass coral bleaching. Nat Clim Chang 8:986–991

Laffoley D., Baxter JM (2016) Explaining ocean warming: causes, scale, effects and consequences. Gland, Switzerland: IUCN.

Layton C, Fulton CJ (2014) Status-dependent foraging behaviour in coral reef wrasses. Coral Reefs 33:345–349

Lejeusne C, Chevaldonné P, Pergent-Martini C, Boudouresque CF, Pérez T (2010) Climate change effects on a miniature ocean: the highly diverse, highly impacted Mediterranean Sea. Trends Ecol Evol 25:250–260

Magalhaes IS, Croft GE, Joyce DA (2013) Altering an extended phenotype reduces intraspecific male aggression and can maintain diversity in cichlid fish. PeerJ 1:e209.

Magalhaes IS, D’Agostino D, Hohenlohe PA, MacColl ADC (2016) The ecology of an adaptive radiation of three-spined stickleback from North Uist, Scotland. Mol Ecol 25:4319–4336

Munday PL, Crawley NE, Nilsson GE (2009) Interacting effects of elevated temperature and ocean acidification on the aerobic performance of coral reef fishes. Mar Ecol Prog Ser 388:235–242

Munday PL, McCormick MI, Nilsson GE (2012) Impact of global warming and rising CO2 levels on coral reef fishes: what hope for the future? J Exp Biol 215:3865–3873

Nagelkerken I, Munday PL (2016) Animal behaviour shapes the ecological effects of ocean acidification and warming: Moving from individual to community-level responses. Glob Chang Biol 22:974–989

Nakamura Y, Feary DA, Kanda M, Yamaoka K (2013) Tropical fishes dominate temperate reef fish communities within western Japan. PLOS One 8:1–8

NASA Goddard space flight center, ocean ecology laboratory, ocean biology processing group. Moderate-resolution imaging spectroradiometer (MODIS) Aqua {Chlorophyll Concentration} Data; NASA OB.DAAC, Greenbelt, MD, USA. Accessed on 29/09/2017

Neuheimer AB, Thresher RE, Lyle JM, Semmens JM (2011) Tolerance limit for fish growth exceeded by warming waters. Nat Clim Chang 1:110–113

Nilsson GE, Crawley N, Lunde IG, Munday PL (2009) Elevated temperature reduces the respiratory scope of coral reef fishes. Glob Chang Biol 15:1405–1412

Nowicki JP, Miller GM, Munday PL (2012) Interactive effects of elevated temperature and CO2 on foraging behavior of juvenile coral reef fish. J Exp Mar Bio Ecol 412:46–51

Pecl GT, Araújo MB, Bell JD, Blanchard J, Bonebrake TC, Chen IC, Clark TD, Colwell RK, Danielsen F, Evengård B, Falconi L, Ferrier S, Frusher S, Garcia RA, Griffis RB, Hobday AJ, Janion-Scheepers C, Jarzyna MA, Jennings S, Lenoir J, Linnetved HI, Martin VY, McCormack PC, McDonald J, Mitchell NJ, Mustonen T, Pandolfi JM, Pettorelli N, Popova E, Robinson SA, Scheffers BR, Shaw JD, Sorte CJB, Strugnell JM, Sunday JM, Tuanmu MN, Vergés A, Villanueva C, Wernberg T, Wapstra E, Williams SE (2017) Biodiversity redistribution under climate change: impacts on ecosystems and human well-being. Science 355:eaai9214

Peng Y, Liu X, Huang G, Wei L, Zhang X (2017) Compensatory growth of juvenile brown flounder Paralichthys olivaceus following low temperature treatment for different periods. J Ocean Univ China 16:326–332

Pörtner HO, Knust R (2007) Climate change affects marine fishes through the oxygen limitation of thermal tolerance. Science 315:95–97

Pörtner HO, Farrell AP (2008) Physiology and climate change. Science 322:690–692

Pörtner HO, Schulte PM, Wood CM, Schiemer F (2010) Niche Dimensions in Fishes: An Integrative View. Physiol Biochem Zool 83:808–826

Pratchett MS, Wilson SK, Munday PL (2015) Effects of climate change on coral reef fishes. In: Camilo Mora (eds) Ecology of Fishes on Coral Reefs. Cambridge University Press 2015, Cambridge, pp 127–134

Pratchett MS, Thompson CA, Hoey AS, Cowman PF, Wilson SK (2018). Effects of coral bleaching and coral loss on the structure and function of reef fish assemblages. In Coral Bleaching (pp. 265–293). Springer, Cham.

Randall JE (1995) Coastal Fishes of Oman. University of Hawaii Press, Honolulu

Richardson LE, Graham NAJ, Pratchett MS, Eurich JG, Hoey AS (2018) Mass coral bleaching causes biotic homogenization of reef fish assemblages. Glob Chang Biol 24:3117–3129

Riegl BM, Purkis SJ (2012) Coral Reefs of the Gulf: Adaptation to Climatic Extremes in the World’s Hottest Sea. In: Riegl B, Purkis S(eds) Coral Reefs of the World, vol 3. Springer, Dordrecht, pp 1–4

Riegl BM, Purkis SJ (2015) Coral population dynamics across consecutive mass mortality events. Glob Chang Biol 21:3995–4005

Riegl BM, Johnston M, Purkis SJ, Howells E, Burt JA, Steiner SCC, Sheppard CRC, Bauman A (2018) Population collapse dynamics in Acropora downingi, an Arabian/Persian Gulf ecosystem-engineering coral, linked to rising temperature. Glob Chang Biol 24:2447– 2462

Rodgers GG, Donelson JM, McCormick MI, Munday PL (2018) In hot water: sustained ocean warming reduces survival of a low-latitude coral reef fish. Mar Biol 165:1–10

Rueda L, Assutí EM, Alvarez-Berastegu D, Hidalgo M (2015) Effect of intra-specific competition, surface chlorophyll and fishing on spatial variation of gadoid’s body condition. Ecosphere 6.10:1–17

Rummer JL, Couturier CS, Stecyk JAW, Gardiner NM, Kinch JP, Nilsson GE, Munday PL (2014) Life on the edge: Thermal optima for aerobic scope of equatorial reef fishes are close to current day temperatures. Glob Chang Biol 20:1055–1066

Saha S et al (2011), updated monthly. NCEP Climate Forecast System Version 2 (CFSv2) Selected Hourly Time-Series Products. Research Data Archive at the National Center for Atmospheric Research, Computational and Information Systems Laboratory. https://doi.org/10.5065/D6N877VB. Accessed 29.09.2017

Scott M, Heupel M, Tobin A, Pratchett M (2017) A large predatory reef fish species moderates feeding and activity patterns in response to seasonal and latitudinal temperature variation. Sci Rep 7:1–9

Sevgili H, Hoşsu B, Emre Y, Kanyilmaz M (2013) Compensatory growth following various time lengths of restricted feeding in rainbow trout (Oncorhynchus mykiss) under summer conditions. J Appl Ichthyol 29:1330–1336

Shinn EA (1976) Coral reef recovery in Florida and the Persian Gulf. Environ Geol 1:241– 254

Shraim R, Dieng MM, Vinu M, Vaughan GO, McParland D, Idaghdour Y, Burt JA (2017) Environmental Extremes Are Associated with Dietary Patterns in Arabian Gulf Reef Fishes. Front Mar Sci 4:285

Sunday JM, Bates AE, Dulvy NK (2011) Global analysis of thermal tolerance and latitude in ectotherms. Proc R Soc B Biol Sci 278:1823–1830

Tewksbury JJ, Huey RB, Deutsch CA (2008) Putting the Heat on Tropical Animals. Science 320:1296–1297

Thornton PK, Ericksen PJ, Herrero M, Challinor AJ (2014) Climate variability and vulnerability to climate change: A review. Glob Chang Biol 20:3313–3328

Tuomainen U, Candolin U (2011) Behavioural responses to human-induced environmental change. Biol Rev 86:640–657

Vaughan GO, Al-Mansoori N, Burt JA (2019) The Arabian Gulf. In Sheppard C(eds) World Seas: an Environmental Evaluation. Academic Press, pp 1–23

Vergés A, Steinberg PD, Hay ME, Poore AGB, Campbell AH, Ballesteros E, Heck KL, Booth DJ, Coleman MA, Feary DA, Figueira W, Langlois T, Marzinelli EM, Mizerek T, Mumby PJ, Nakamura Y, Roughan M, van Sebille E, Gupta A Sen, Smale DA, Tomas F, Wernberg T, Wilson SK (2014) The tropicalization of temperate marine ecosystems: Climate-mediated changes in herbivory and community phase shifts. Proc R Soc B Biol Sci 281:

Wernberg T, Smale DA, Tuya F, Thomsen MS, Langlois TJ, De Bettignies T, Bennett S, Rousseaux CS (2013) An extreme climatic event alters marine ecosystem structure in a global biodiversity hotspot. Nat Clim Chang 3:78–82

Wilkinson CR (1999) Global and local threats to coral reef functioning and existence: Review and predictions. Mar Freshw Res 50:867–878

Wong BBM, Candolin U (2015) Behavioral responses to changing environments. Behav Ecol 26:665–673

Zhou L, Zeng L, Fu D, Xu P, Zeng S, Tang Q, Chen Q, Chen L, Li G (2016) Fish density increases from the upper to lower parts of the Pearl River Delta, China, and is influenced by tide, chlorophyll-a, water transparency, and water depth. Aquat Ecol 50:59–74

